# Development of an *in vivo* electroporation-based chromosomal engineering technique in the murine uterine epithelium

**DOI:** 10.1101/2025.02.25.640089

**Authors:** Satoru Iwata, Yumi Miura, Takashi Iwamoto

## Abstract

Clustered regularly interspaced palindromic repeat (CRISPR)/CRISPR-associated protein (Cas)-based *in vivo* chromosomal rearrangements are a promising approach for generating model organisms with specific chromosomal abnormalities. However, conventional *in vivo* methods rely on viral vectors, which are expensive, require specialized equipment, and pose potential safety risks, thereby limiting their widespread application. To overcome the limitations above, we developed a novel, efficient, and cost-effective *in vivo* chromosomal engineering strategy using CRISPR ribonucleoprotein electroporation for the murine uterine epithelium. Our method successfully induced translocations at multiple loci and repaired a 57.8-Mb inversion. The findings of the present study establish *in vivo* electroporation as a practical alternative to traditional chromosomal engineering methods and provide a foundation for its broader application in genome editing technologies.

## Introduction

Chromosomal abnormalities, particularly inversions and translocations, play a crucial role in the development of various cancers and hereditary disorders. Chromosomal rearrangements not only alter gene expression and promote tumorigenesis by dysregulating oncogenes and tumor suppressor genes (Mitelman et al., 2007; Chen et al., 2010) but also induce mutations in genes involved in essential physiological functions, thereby causing infertility and various hereditary diseases (Lee et al., 2006; Feuk, 2010; Pellestor et al., 2011; Zepeda-Mendoza et al., 2019). The precise and efficient repair of such chromosomal abnormalities is essential for treatment, and the clustered regularly interspaced palindromic repeat (CRISPR)/CRISPR-associated protein (Cas) system is widely used for this process (Katti et al., 2022; Nambiar et al., 2022).

The CRISPR/Cas system has been used successfully to generate model organisms with specific chromosomal abnormalities (Blasco et al., 2014; Maddalo et al., 2014; Reimer et al., 2017). These models have contributed significantly to the elucidation of various disease mechanisms and the development of novel therapeutic strategies. Furthermore, newly developed prime editing techniques, such as WT-PE and PETI, exhibit high precision in inducing inversions and translocations in cultured human cells (Tao et al., 2022; Kweon et al., 2023). These techniques have great potential for the treatment of diseases caused by chromosomal rearrangements.

However, *in vivo* approaches of inducing chromosomal rearrangements using the CRISPR/Cas system often require a combination of viral vectors, which entails considerable effort, cost, specialized experimental facilities, and potential safety risks, thereby limiting their widespread application. Moreover, to the best of our knowledge, no study has demonstrated successful induction of chromosomal rearrangements via prime editing *in vivo* to date. Therefore, the development of simple, safe, and efficient *in vivo* chromosomal rearrangement techniques is necessary to treat various genetic disorders and cancers.

We have previously performed *in vivo* electroporation using RecQ-like helicase 5 (*Recql5*) mutations to efficiently induce complex chromosomal rearrangements (CCRs) by delivering CRISPR ribonucleoproteins (RNPs) into fertilized mouse embryos in the oviduct (Iwata et al., 2024). Nevertheless, the widespread application of our approach is limited by restrictions on the use of fertilized embryos and ethical regulations surrounding human embryo editing (Greely, 2019). In addition, our *Recql5*-based approach revealed the involvement of replication-based complex rearrangement mechanisms, such as chromoanasynthesis-like events, in the repair of double-strand breaks (DSBs) associated with chromosomal rearrangements. Chromoanasynthesis is characterized by extensive structural alterations, including localized duplications and triplications, caused by replication fork stalling, template switching (FoSTeS), and microhomology-mediated break-induced replication (MMBIR) (Pellestor & Gatinois, 2018). Such mechanisms increase the risk of unintended genomic alterations, thereby limiting their application in chromosomal engineering.

In this study, we extended the scope of chromosomal engineering from fertilized embryos to organ tissues using *in vivo* electroporation. Specifically, we modified the CRISPR RNP electroporation method developed by Kobayashi et al. (2023) and assessed its safety and efficiency in inducing chromosomal rearrangements in the murine uterine epithelium. We also examined the presence of FoSTeS/MMBIR during chromosomal repair. Gynecological cancers, particularly uterine cancer, are among the most frequently diagnosed malignancies worldwide, with chromosomal abnormalities such as translocations and inversions being common hallmark features of tumorigenesis (Lu et al., 2021). The present study demonstrates the feasibility of *in vivo* electroporation as a practical and efficient method for inducing chromosomal rearrangements in the uterine epithelium.

## Results

### Establishment of an *in vivo* editing technique for CCR in the mouse uterine epithelium

Several clinical trials have demonstrated the potential of CRISPR-based genome editing in the treatment of various genetic disorders and cancers (Morshedzadeh et al., 2024). However, their potential for *in vivo* chromosomal rearrangements remains to be elucidated. To address this, we modified the CRISPR RNP electroporation method developed by Kobayashi et al. (2023) and applied it to induce chromosomal rearrangements in the murine uterine epithelium (Figure 1A; Figure S1). CRISPR RNPs were first injected into the uterine lumen via microinjection and subsequently introduced into uterine epithelial cells through electroporation. The use of a microinjector allowed for more precise and controlled reagent delivery, reducing mechanical stress on the tissue and minimizing leakage, which is expected to facilitate a more localized and consistent distribution of RNPs within the uterine lumen.

**Figure 1.**
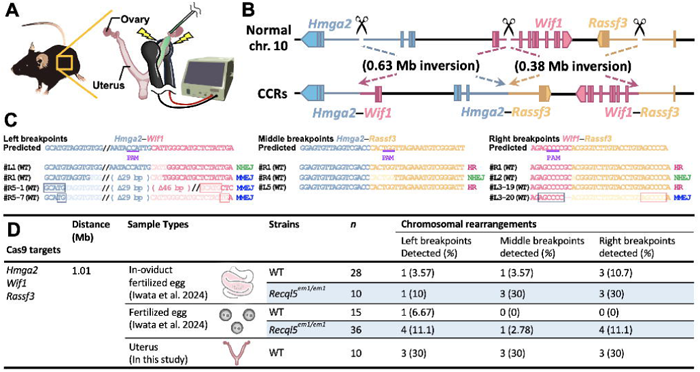
CCRs in the uterine epithelium induced via *in vivo* electroporation. (A) Schematic overview of the experimental procedure for CCR induction by *in vivo* electroporation. (B) Diagram illustrating the CCRs among *Hmga2*, *Wif1*, and *Rassf3* on chromosome 10. (C) Sequence alignment of PCR products corresponding to the genomic breakpoint junctions, *Hmga2–Wif1*, *Hmga2–Rassf3*, and *Wif1–Rassf3*. Boxed regions indicate the microhomology sequences. (D) Comparative analysis of CCR induction efficiency via *in vivo* electroporation of within-oviduct fertilized embryos, fertilized embryos, and uterine epithelium.

As an initial trial, we tested 1.1-Mb CCRs within mouse chromosome 10, encompassing the *Hmga2*, *Wif1*, and *Rassf3*, which are implicated in various human cancers and are mapped in a configuration similar to that of human chromosome 12 (Pareja et al., 2019) (Figure 1B). This region was previously targeted in fertilized embryos (Iwata et al., 2024), and was selected for gene editing in somatic tissues to enable direct comparison of editing efficiencies. On postoperative day 7, genomic DNA was extracted from the tissue and the achievement of the intended chromosomal rearrangements was assessed via PCR and Sanger sequencing.

Sequencing of the genomic DNA isolated from the uterus revealed that chromosomal rearrangements were successfully induced at three different breakpoints (Figure 1C). Sequence analysis confirmed that HR, nonhomologous end-joining (NHEJ), and microhomology-mediated end-joining (MMEJ) were involved in breakpoint repair (Figure 1C). Although the results suggest relatively high occurrence of genome-editing events compared to previous reports in fertilized embryos (Figure 1D; Iwata et al., 2024), direct comparisons should be interpreted with caution because of the intrinsic differences between embryonic and somatic tissues. In fertilized embryos, genome editing occurs at the single-cell stage, enabling precise determination of the proportion of successfully edited embryos. In contrast, somatic tissues consist of a mosaic population of edited and unedited cells, making absolute quantification of editing efficiency more complex. Nevertheless, consistent detection of chromosomal rearrangements at multiple breakpoints suggests that *in vivo* electroporation effectively facilitates genome editing in somatic tissues.

### Efficient induction of interchromosomal translocations in the mouse uterine epithelium

Next, we evaluated the feasibility of inducing translocations, in which segments from the two chromosomes were exchanged. The functional intergenic repeating RNA element (*Firre*) gene located on the X chromosome was selected as the target locus. The *Firre* locus plays a central role in interchromosomal interactions by facilitating gene activation through three-dimensional spatial interactions with regulatory regions on other chromosomes (Maass et al., 2019) (Figure 2A). Therefore, *Firre* serves as a key hub for transcriptional regulation and is expected to provide a structural basis for efficient translocation induction. We designed experiments to induce translocations between the *Eef1a1* neighborhood locus (*Eef1a1N*) on chromosome 9 and the *Atf4* neighborhood locus (*Atf4N*) on chromosome 15, both of which interact with the *Firre* loci. Additionally, we targeted the translocations between *Ypel4* neighborhood locus (*Ypel4N*) on chromosome 2 and *Atf4N* on chromosome 15, as these loci are also physically linked to the *Firre* locus (Figure 2A, B). On postoperative day 7, the predicted translocations were detected in uterine samples (Figure 2C, D). The observed translocation efficiencies were 33.3 and 66.7% for *Eef1a1N–Atf4N* and 40 and 30% for *Ypel4N–Atf4N*, respectively (Figure 2E). These findings suggest a high efficiency of translocation induction in this experimental setting and support the potential effectiveness of this *in vivo* electroporation-based approach.

**Figure 2.**
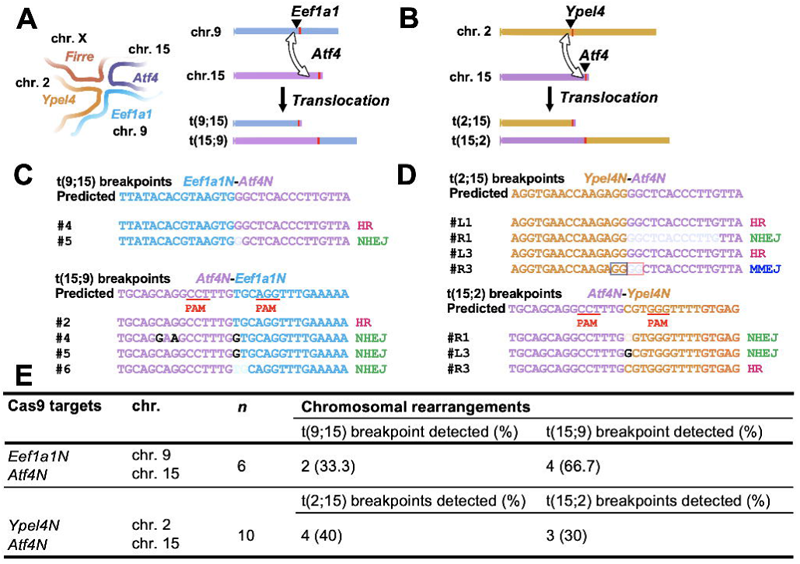
Translocations at the *Firre* locus in the uterine epithelium induced via *in vivo* electroporation. (A) Schematic representation of the translocation between the *Eef1a1N* on chromosome 9 and *Atf4N* on chromosome 15. (B) Sequence alignment of the PCR products corresponding to the genomic breakpoint junctions of *Eef1a1N* and *Atf4N*. (C) Schematic representation of the translocation between the *Ypel4N* on chromosome 2 and *Atf4N* on chromosome 15. (D) Sequence alignment of the PCR products corresponding to the genomic breakpoint junctions of *Ypel4N* and *Atf4N*. Boxed regions indicate the microhomology sequences. (E) Overall translocation efficiency in the uterine epithelium.

### Repair of the large-scale 57.8-Mb chromosomal inversion, *In(6)1J*, in the mouse uterine epithelium

CRISPR/Cas9 technology and its derivatives, including base and prime editors, offer new therapeutic avenues for cancer and genetic disorders (Porto et al., 2020; Katti et al., 2022; Chen et al., 2023). Here, we explored the feasibility of repairing chromosomal abnormalities in the mouse uterine epithelium via CRISPR/Cas9-based *in vivo* electroporation. We aimed to repair the large-scale 57.8-Mb inversion *In(6)1J* on chromosome 6, which was first identified in the C3H/HeJ strain in 2006 (Akeson et al., 2006; Ackert-Bicknell et al., 2007) and whose breakpoints were previously mapped in 2021 (Iwata et al., 2021). This inversion spans approximately 40% of the total length of chromosome 6, and is implicated in specific phenotypes, including low bone mass and increased bone marrow adiposity (Ackert-Bicknell et al., 2007). In the present study, we designed gRNAs near breakpoints and employed a strategy to repair chromosomal inversions using ssODNs corresponding to normal genomic sequences (Figure 3A). Repair efficiencies of 18.2 and 36.4% were observed at the left and right breakpoints, respectively (Figure 3B, C). The successful repair of this 57.8-Mb inversion demonstrated that this *in vivo* approach is applicable to the repair of large-scale chromosomal abnormalities. To the best of our knowledge, this is the largest chromosomal rearrangement targeted for genome editing.

**Figure 3.**
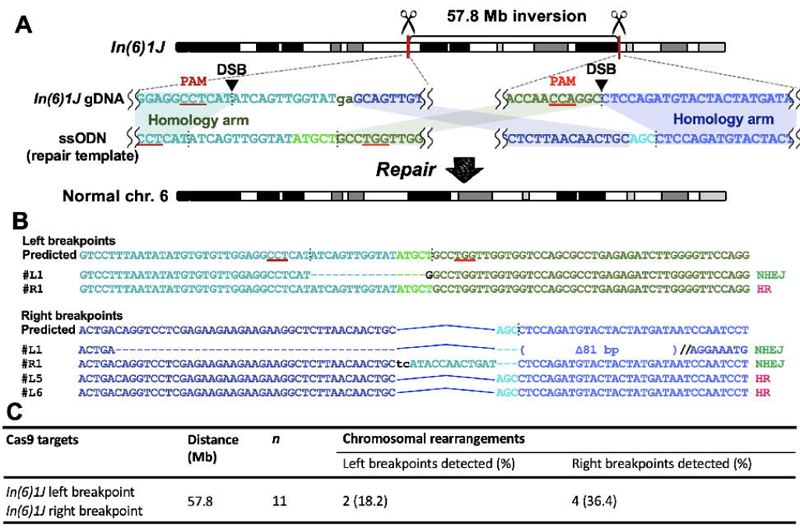
Repair of a large-scale chromosomal inversion in the uterine epithelium via *in vivo* electroporation. (A) Schematic representation of the CRISPR/Cas9-based strategy to repair the 57.8-Mb inversion, *In(6)1J*, on chromosome 6. (B) Sequence alignment of the PCR products corresponding to the left and right breakpoint junctions after repair. (C) Overall repair efficiency in the uterine epithelium.

### *In vivo* chromosomal engineering of the mouse uterine epithelium by targeting multiple genetic loci

Based on these findings, a broad investigation of locus-dependent differences in editing efficiency is crucial. Therefore, we evaluated the ability of *in vivo* electroporation to induce translocations and inversions at other genetic loci. To introduce oncogenic fusion genes, translocations were induced between *Ncoa2* and *Greb1*, as well as between *Ywhae* and *Nutm2*, both of which are associated with uterine sarcomas (Figure 4A, B). GREB1–NCOA2 fusion integrates the estrogen-responsive growth-promoting function of GREB1 with the nuclear receptor coactivator function of NCOA2, thereby enhancing hormone-responsive cancer cell proliferation, tumor initiation, and progression (Brunetti et al., 2018). YWHAE–NUTM2 fusion promotes oncogenesis by inducing cyclin D1 overexpression via the regulation of the RAF/mitogen-activated protein kinase and Hippo signaling pathways, driving the proliferation of high-grade endometrial stromal sarcoma cells (Ou et al., 2021). Additionally, we re-examined the 7.67-Mb inversion previously reported for fertilized embryos (Iwata et al., 2024) (Figure S2A). This targeted locus exhibits FoSTeS/MMBIR in fertilized embryos carrying *Recql5* mutations (Iwata et al., 2024).

**Figure 4.**
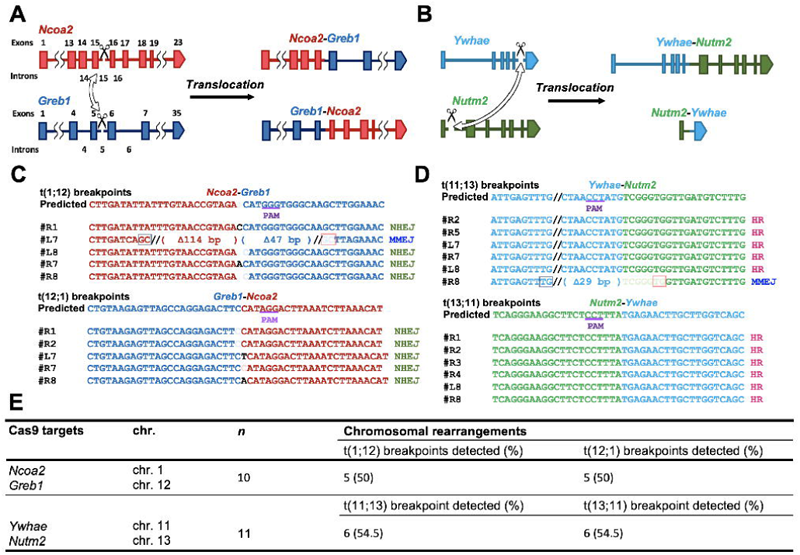
Oncogenic translocations in the uterine epithelium induced via *in vivo* electroporation. (A) Schematic representation of the translocation between the *Ncoa2* on chromosome 1 and *Greb1* on chromosome 12. (B) Sequence alignment of the PCR products corresponding to the genomic breakpoint junctions of *Ncoa2* and *Greb1*. Boxed regions indicate the microhomology sequences. (C) Schematic representation of the translocation between *Ywhae* on chromosome 11 and *Nutm2* on chromosome 13. (D) Sequence alignment of the PCR products corresponding to the genomic breakpoint junctions of *Ywhae* and *Nutm2*. (E) Overall translocation efficiency in the uterine epithelium.

Successful chromosomal engineering was observed at all the tested genetic loci (Figure 4C, D, E; Figure S2B, C). Notably, no FoSTeS/MMBIR-associated events were detected at any of the loci analyzed in this study. Although the fusion genes were successfully induced during the translocations, no tumors were detected during the observation period. Given that tumor development often requires an extended timeframe (Klein, 2009), further long-term investigations are required. Collectively, our findings suggest that *in vivo* electroporation effectively induced chromosomal rearrangements at multiple genetic loci.

## Discussion

Using CRISPR RNP electroporation, we successfully induced interchromosomal translocations and repaired a large-scale 57.8-Mb inversion, demonstrating the feasibility of this approach for correcting chromosomal abnormalities. Given that large-scale chromosomal rearrangements are frequently associated with genetic disorders and cancers (Popova et al., 2012; Pellestor, 2019; Comaills et al., 2023), the ability to efficiently correct them *in vivo* presents a promising avenue for therapeutic genome editing.

Previous studies, such as those by Reimer et al. (2017), utilized lentiviral vectors to induce chromosomal translocations, including the t(11, 19)/MLL-ENL translocation. While effective, lentiviral-based methods pose risks, such as insertional mutagenesis and prolonged Cas9 expression, increasing the likelihood of off-target effects and genomic instability (Bushman, 2020). Additionally, their production is costly, complex, and requires biosafety containment. Adeno-associated virus-based genome editing has been widely adopted because of its low immunogenicity and stable transgene expression (Verdera et al., 2020). However, its small packaging capacity (∼4.7 kb) restricts the delivery of large genome-editing components such as Cas9 and multiple guide RNAs. Adenoviral vectors offer an alternative viral delivery system with high packaging capacity (∼36 kb) and robust transgene expression. Unlike lentiviral vectors, AdVs do not integrate into the host genome, thereby minimizing the risk of insertional mutagenesis (Li and Lieber, 2019). Nevertheless, their high immunogenicity often triggers strong inflammatory responses, leading to reduced efficacy *in vivo* (Asmamaw Mengstie, 2022). In contrast, our CRISPR RNP electroporation approach eliminates these risks by enabling transient genome editing, thereby reducing the potential for genomic instability. Additionally, electroporation does not require viral vector production, simplifies the experimental workflow, and significantly reduces costs. These advantages highlight the safety and practicality of *in vivo* electroporation for chromosomal engineering, particularly for applications requiring precise genome editing in somatic tissues.

In addition to these advantages, a key observation in this study was that targeted chromosomal rearrangements were successfully induced and consistently detected in somatic tissues at a higher frequency than in fertilized embryos. To the best of our knowledge, CRISPR RNP electroporation has not been reported to successfully induce translocation in fertilized embryos. This discrepancy may be attributed to several factors, including differences in the number of available target cells, cell cycle duration, and DNA repair mechanisms, which are predominant in each cellular environment. Unlike fertilized embryos, in which genome editing occurs at the one-cell stage, somatic tissues consist of numerous target cells, increasing the absolute number of successfully edited cells and enabling the consistent detection of chromosomal rearrangements. However, in fertilized embryos, a rapid cell cycle may constrain the time window available for DNA repair (Khokhlova et al., 2020), particularly in the context of large-scale chromosomal rearrangements. These rearrangements can induce genomic instability, reduce cell viability, and limit the efficiency of chromosomal engineering. Previous studies have reported that the efficiency of CRISPR-based genome editing is significantly influenced by the cell cycle phase (Abe et al., 2020), and that rapid embryonic cell division may leave insufficient time for precise repair. In contrast, the prolonged cell cycle in somatic tissues provides more time for accurate DNA repair (Kermi et al., 2019), which may have facilitated the successful induction of translocation in the present study. Additionally, uterine epithelial cells, particularly those in the metestrus phase, exhibit more stable cell cycle regulation (Baranda-Avila et al., 2009), potentially providing a favorable environment that supports efficient chromosomal engineering while mitigating the risk of genomic instability-induced cell death.

Overall, the findings highlight the feasibility of CRISPR RNP-based *in vivo* electroporation as a viable approach for chromosomal engineering of somatic tissues. This approach offers a transient and flexible alternative to virus-based methods, thereby avoiding the concerns related to stable genome integration. The successful repair of a 57.8-Mb chromosomal inversion further underscores its potential for large-scale genome correction, offering new opportunities in cancer research, congenital disorder treatment, and regenerative medicine. Further optimization and long-term validation studies are essential to fully utilize this approach for therapeutic applications.

## Limitations

In the present study, chromosomal rearrangements were efficiently induced in uterine epithelial samples, although variability in genome-editing efficiency was observed. Therefore, further optimization of the electroporation parameters, including voltage and pulse duration, is necessary to improve the effectiveness of this technique. In addition, its feasibility in other tissues and cell types should be explored to enhance its applicability.

Future studies with extended observation periods and comprehensive biological evaluations, including tumorigenic risk assessments, are required, to determine the long-term biological effects of this method. This study provides important insights into the feasibility of *in vivo* electroporation-based chromosomal engineering. However, further rigorous validation is essential to establish its safety and efficacy for future clinical applications.

## Materials and methods

### Experimental animals

Wild-type (WT) mice (C57BL/6NCrSlc; Japan SLC, Shizuoka, Japan) and *In(6)1J* inversion-carrying mice (C3H/HeJJcl; CLEA Japan, Tokyo, Japan) were used in this study. Sexually mature female mice aged 8 weeks or older were selected for experiments. All animals were maintained under controlled environmental conditions at a constant temperature of 22 ± 2 °C and humidity of 50 ± 10% under a 12/12-h light/dark cycle. All animal experiments were approved by the Institutional Animal Care and Use Committee of Chubu University (permit number #202410012; Kasugai, Aichi, Japan) and adhered to the guidelines.

### CRISPR RNP and single-stranded oligodeoxynucleotide (ssODN) preparation

CRISPR guide RNAs (gRNAs) were designed using CHOPCHOP (http://chopchop.cbu.uib.no/; Table S1) (Labun et al., 2019). The CRISPR RNP complex was assembled using Alt-R S.p. Cas9 Nuclease 3NLS (Integrated DNA Technologies, Coralville, IA, USA) and a custom-designed gRNA consisting of a CRISPR RNA (crRNA):trans-activating CRISPR RNA (tracrRNA) duplex (Integrated DNA Technologies). After resuspending in the nuclease-free duplex buffer (Integrated DNA Technologies) at a final concentration of 4,000 ng/μL, crRNA and tracrRNA were mixed at equimolar ratios, heated at 95 °C for 10 min, and gradually cooled to 25 °C to facilitate duplex formation. Then, the crRNA:tracrRNA duplex was incubated with Alt-R S.p. Cas9 Nuclease 3NLS at 25 °C for 10 min to assemble the RNP complex. Specific concentrations are indicated in the following section: *in vivo* genome editing of the murine endometrium. Subsequently, ssODNs were synthesized by Eurofins Genomics (Tokyo, Japan) to facilitate the seamless joining of the two DNA sequences, ensuring that the junction was positioned at the center of the predicted cleavage sites located within 3 bp of the protospacer adjacent motif (PAM) sequences (Table S2). Moreover, 5′ and 3′ ends of ssODNs were modified with two consecutive phosphorothioate (*) linkages to enhance the HR efficiency (Renaud et al., 2016) (Table S2).

### *In vivo* genome editing of the murine endometrium

Female metestrus or diestrus mice were subjected to *in vivo* genome editing of the murine endometrium using a modified version of the method previously described by Kobayashi et al. (2023). To generate complex CCRs, the following CRISPR solution was used: 520 ng/μL Alt-R S.p. Cas9 Nuclease 3NLS, 30 μM crRNA:tracrRNA duplex each for the left, middle, and right targets, 158 ng/μL ssODN each for the left, middle, and right targets, and 0.02% Fast Green FCF (Wako, Osaka, Japan) marker diluted in the Opti-MEM (Thermo Fisher Scientific, Waltham, MA, USA). To generate chromosomal inversions and translocations, the following CRISPR solution was used: 520 ng/μL Alt-R S.p. Cas9 Nuclease 3NLS, 30 μM crRNA:tracrRNA duplex each for the left and right targets in inversions and for each breakpoint on the respective chromosomes in translocations, 158 ng/μL ssODN each for the left and right targets in inversions and for each breakpoint on the respective chromosomes in translocations, and 0.02% Fast Green FCF (Wako) marker diluted in the Opti-MEM (Thermo Fisher Scientific). Prior to electroporation, the female mice were anesthetized with a mixture of medetomidine (0.75 mg/kg, Nippon Zenyaku Kogyo, Fukushima, Japan), midazolam (4 mg/kg, Sandoz, Tokyo, Japan), and butorphanol (5 mg/kg, Meiji Seika Pharma, Tokyo, Japan).

Glass micropipettes were first pulled using a vertical capillary puller (NARISHIGE, Tokyo, Japan) to create fine-tipped capillaries suitable for microinjection. Before use, the tip of each capillary was cut diagonally to a length of 0.5 mm using micro-scissors to facilitate smooth reagent flow. The cut capillary was then connected to a syringe via a silicone tube and filled with the CRISPR solution, ensuring complete reagent loading. Once filled, the capillary was attached to a capillary holder and secured in place using a flexible stand. The CRISPR solution (2 μL) was then introduced into the uterus via microinjection from the downstream part of the oviduct using the prepared glass micropipette and the FemtoJet 4i microinjector (Eppendorf, Hamburg, Germany) under the following parameters: injection pressure (pi) = 100 hPa, injection duration (ti) = 0.2 s, and compensation pressure (pc) = 0 hPa. To prevent reagent backflow, the oviduct near the ovary was clamped using a hemostatic clip (Natsume Seisakusho, Tokyo, Japan). The uterus was clamped to the cervical side using a hemostatic clip (Natsume Seisakusho) to prevent leakage. Following the injection, the uterine horns were clamped using tweezer electrodes (LF650P5; BEX, Tokyo, Japan), and electroporation was performed using a CUY21EDIT II electroporator (BEX). The electrode orientation was adjusted and three additional sets of electric pulses were applied. The following electroporation parameters were used: primary large pulse (PLP): 20 V, 30.0-ms pulse duration, 50.0-ms pulse interval, 3 pulses, 10% decay (± pulse orientation). Secondary decay pulse (Pd): 10 V, 50.0-ms pulse duration, 50.0-ms pulse interval, 3 pulses, 40% decay (± pulse orientation). After electroporation, the uterine horns were repositioned and the incisions were sutured. To reverse the effects of medetomidine, atipamezole hydrochloride (0.75 mg/kg, Nippon Zenyaku Kogyo, Fukushima, Japan) was intraperitoneally administered postoperatively.

### Analysis of the CRISPR/Cas9-engineered mice

To screen for CRISPR/Cas9-induced mutations, genomic DNA was extracted from the uterine horns of founder mice using a DNeasy Blood & Tissue Kit (Qiagen, Hilden, Germany) and subjected to polymerase chain reaction (PCR) amplification using EmeraldAmp PCR Master Mix (Takara Bio, Shiga, Japan). For nested PCR, the first round consisted of 40 cycles of denaturation at 98 °C for 10 s, annealing at 60 °C for 30 s, and extension at 72 °C for 1 min, followed by a final indefinite hold at 10 °C. The second round of PCR consisted of 15 cycles of denaturation at 98 °C for 10 s, annealing at 63 °C for 30 s, and extension at 72 °C for 30 s, with a final indefinite hold at 10 °C. The resulting PCR products were purified using a NucleoSpin Gel and PCR Cleanup Kit (Takara Bio) and cloned into the pTAC-1 vector (Biodynamics, Tokyo, Japan). Individual clone sequences were determined using Sanger sequencing (Eurofins Genomics). All PCR primers used for genotyping are listed in Table S3.

## Acknowledgments

The authors thank the laboratory members for their support with animal care and the experiments. The authors would like to thank Editage (www.editage.com) for English language editing.

## Author contributions

SI designed and performed the experiments and drafted the manuscript. YM performed animal experiments. TI supervised the study and revised the manuscript. All the authors have read and approved the final manuscript.

