## SUPPLEMENTARY INFORMATION for "Development of an *in vivo* electroporation-based chromosomal engineering technique in the murine uterine epithelium"

**Supplementary Figures:**
**Figure S1:** Experimental procedure for the induction of chromosomal rearrangements in the mouse uterine epithelium via *in vivo* electroporation.
**Figure S2 :** Inversion in the uterine epithelium induced via *in vivo* electroporation.

**Supplementary Tables:**
**Table S1:** gRNAs used in the present study.
**Table S2:** Single-stranded oligodeoxynucleotides (ssODNs) used in the present study.
**Table S3:** Primers used in the present study.

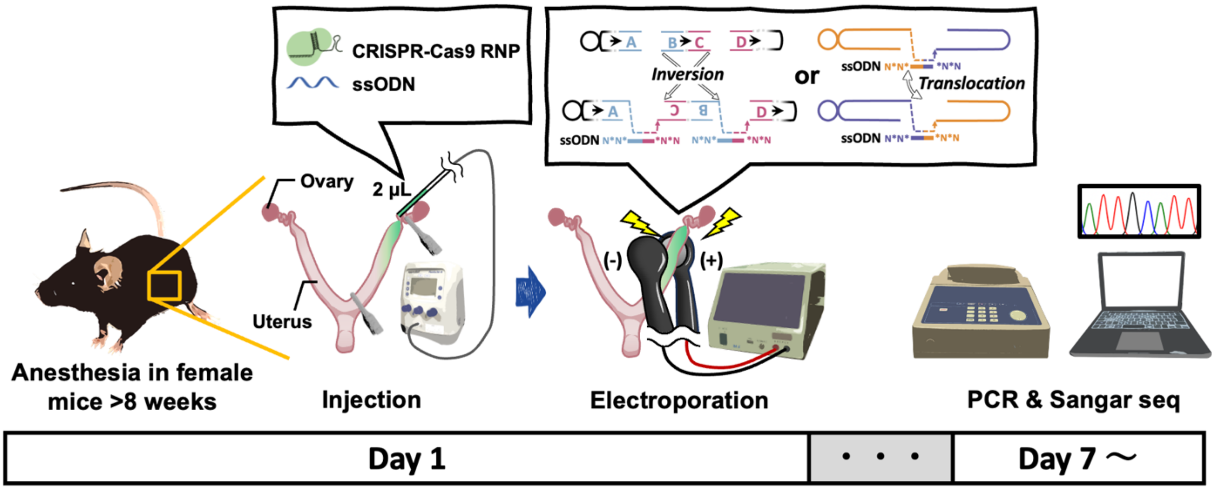


**Figure S1.**
**Experimental procedure for the induction of chromosomal rearrangements in the mouse uterine epithelium via *in vivo* electroporation.**

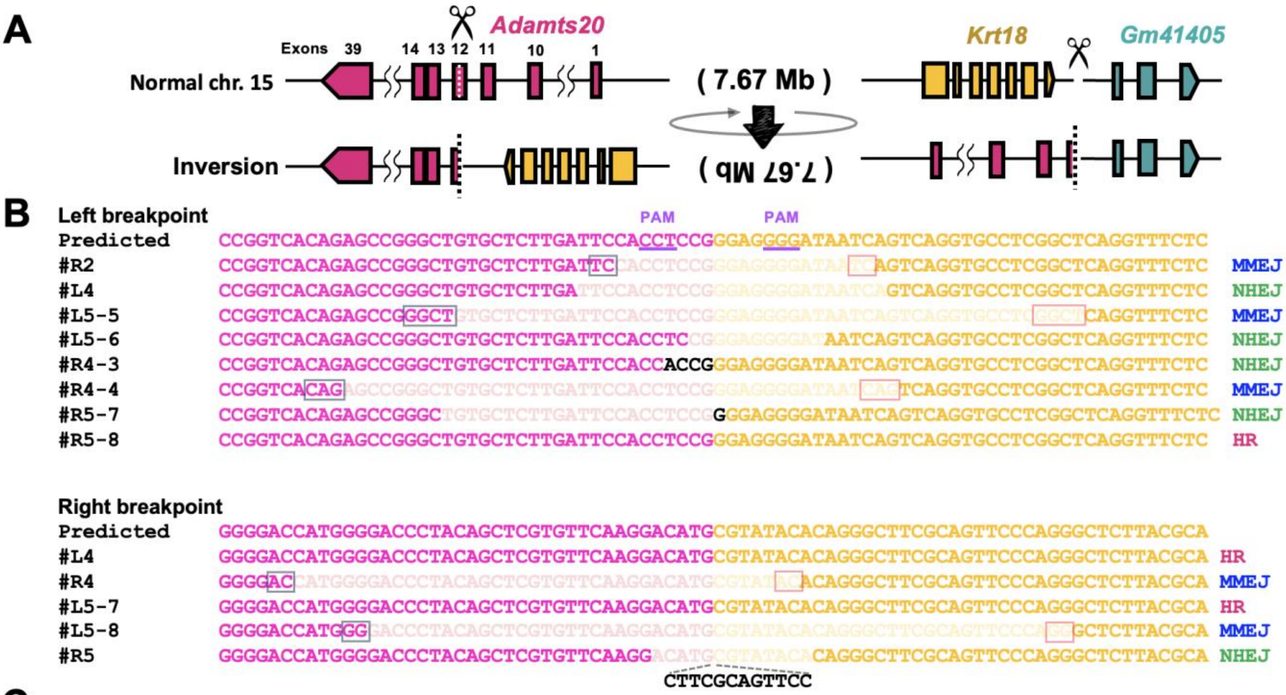


**C**

| Cas9 targets | Distance (Mb) | *n* | Chromosomal rearrangements | |
|---|---|---|---|---|
| | | | Left breakpoints detected (%) | Right breakpoints detected (%) |
| *Adamts20* *K18N* | 7.67 | 10 | 5 (50) | 4 (40) |

**Figure S2.**

**Inversion in the uterine epithelium induced via *in vivo* electroporation.**

(A) Schematic representation of the chromosomal rearrangement caused by the inversion between *Adamts20* and *Krt18* neighborhood locus (*K18N*) on chromosome 15. (B) Sequence alignment of the PCR products corresponding to the genomic breakpoint junctions of *Adamts20* and *K18N* locus. Boxed regions indicate the microhomology sequences. (C) Overall inversion efficiency in the uterine epithelium.

### Table S1: gRNAs used in the present study.

| Target loci | Genomic location (mm10/GRCm38) | Target Sequences (PAM) | CHOPCHOP (https://chopchop.cbu.uib.no/) | | | | |
|---|---|---|---|---|---|---|---|
| | | | Number of mismatches | | | | Efficiency |
| | | | 0 | 1 | 2 | 3 | |
| *Hmga2* | chr10: 120416213 | GGAGTGTTAGGTCGACCCAA (TGG) | 0 | 0 | 0 | 0 | 67.29 |
| *Wif1* | chr10: 121049539 | ATAGAGCATGCCCAATGGCG (GGG) | 0 | 0 | 0 | 0 | 60.84 |
| *Rassf3* | chr10: 121431836 | ACAGGTACAAGACCCGTCAC (TGG) | 0 | 0 | 0 | 2 | 59.01 |
| *Eef1a1N* | chr9: 78477382 | ACTTATACACGTAAGTGTGC (AGG) | 0 | 0 | 0 | 2 | 55.34 |
| *Atf4N* | chr15: 80258539 | CGTAACAAGGGTGAGCCCAA (AGG) | 0 | 0 | 0 | 2 | 64.88 |
| *Ypel4N* | chr2: 84738819 | CCAGGTGAACCAAGAGGCGT (GGG) | 0 | 0 | 0 | 5 | 59.00 |
| *In(6)1J* left breakpoint | chr6: 63000847 | TGCtcATACCAACTGATATG (AGG) | - | | | | |
| *In(6)1J* right breakpoint | chr6: 120827195 | TAGTAGTACATCTGGAGGCC (TGG) | - | | | | |
| *Ywhae* | chr11: 75760846 | GACCAAGCAAGTTCTCACAT (AGG) | 0 | 0 | 0 | 5 | 70.57 |
| *Nutm2* | chr13: 50467977 | AGACATCAACCACCCGATAA (AGG) | 0 | 0 | 0 | 3 | 51.94 |
| *Ncoa2* | chr1: 13161287 | TTATTTGTAACCGTAGACAT (AGG) | 0 | 0 | 0 | 7 | 65.93 |
| *Greb1* | chr12: 16734411 | GTTAGCCAGGAGACTTCCAT (GGG) | 0 | 0 | 0 | 5 | 62.57 |
| *Adamts20* | chr15: 94347708 | TCGTGTTCAAGGACATGCGG (AGG) | 0 | 0 | 1 | 3 | 71.44 |
| *K18N* | chr15: 102034830 | GAAGCCCTGTGTATACGGGA (GGG) | 0 | 0 | 1 | 4 | 61.49 |

### Table S2: Single-stranded oligodeoxynucleotides (ssODNs) used in the present study.

| Target loci | Target Sequences (5′–3′) |
|---|---|
| *Hmga2*–*Wif1* (Left) | PAM<br>C*G*TTGCAGCATGTAGGTGTGGTAGCATCTGCACCCACCAAAATACCATTG<br>CATTGGGCATGCTCTATTGAAGGCTGAGTCCAGAGCTCTCCATTGGGC*A*T |
| *Hmga2*–*Rassf3* (Middle) | G*A*AGCACGAAGAAAAGAGTTTAAAAATTCAAAGGGAGTGTTAGGTCGACC<br>CACTGGTTAGAAATGTCGGGATTGGTTGCCCCCTGCCTCCTAGGCACT*G*G |
| *Wif1*–*Rassf3* (Right) | T*A*ACCCAAAAAAGATTTTTTTAAAAAACCACGCCTTACTGAGAGCCCCGC<br>ACGGGTCTTGTACCTGTAGCCCCATGGGCCATGGGTTCTTGGCTAATT*T*T |
| *Eef1a1N*–*Atf4N* t(9:15) | T*G*TCTTCAAAGCACCAGTAGATGGTGCTACTCCACTTATACACGTAAGTG<br>GGCTCACCCTTGTTACGCACAGAAGCTAGGCTGTAAGTAGTTAAGTCT*C*T |
| *Atf4N*–*Eef1a1N* t(15:9) | G*G*TGTGGGTAGGATGATACAGCAGCCTCCCACTTCTGCAGCAGGCCTTTG<br>TGCAGGTTTGAAAAACGGTGTTTGTATCCAGCAGACACTTGCTATCAA*T*T |
| *Ypel4N*–*Atf4N* t(2:15) | T*C*CTCCTCCACCTGTTTCACAAGCGGTAAGCATCCAGGTGAACCAAGAGG<br>GGCTCACCCTTGTTACGCACAGAAGCTAGGCTGTAAGTAGTTAAGTCT*C*T |
| *Atf4N*–*Ypel4N* t(15:2) | G*G*TGTGGGTAGGATGATACAGCAGCCTCCCACTTCTGCAGCAGGCCTTTG<br>CGTGGGTTTTGTGAGCTTGTCCAAAGGGAAGGAATATCTAAGTGTCTT*C*A |
| *In(6)1J* left breakpoint | C*T*GGAAGTCCTTTAATATATGTGTGTTGGAGGCCTCATATCAGTTGGTAT<br>ATGCTGCCTGGTTGGTGGTCCAGCGCCTGAGAGATCTTGGGGTTCCAGGTTA<br>A*T*T |
| *In(6)1J* right breakpoint | C*T*GGGAACTGACAGGTCCTCGAGAAGAAGAAGAAGGCTCTTAACAACTGC<br>AGCCTCCAGATGTACTACTATGATAATCCAATCCTTAGTAGAGAATAACAG*<br>G*A |
| *Ncoa2*–*Greb1* t(1:12) | A*A*GATCAGTTGCCTATACAGCTCCTACTTGATATTATTTGTAACCGTAGA<br>CATGGGTGGGCAAGCTTGGAAACTGCAGAGGAAGCAGTGGCACAAAGG*C*T |
| *Greb1*–*Ncoa2* t(12:1) | G*T*CCTTCCTCTGGGTTTTAGATGATCTGTAAGAGTTAGCCAGGAGACTTC<br>CATAGGACTTAAATCTTAAACATTAGCTCAATTCTAAAAAGTCTGTGT*G*T |
| *Ywhae*–*Nutm2* t(11:13) | T*T*TCTGGCAAAATTGAGTTTGATTTTTTACTTTATTTTTCCTAACCTATG<br>TCGGGTGGTTGATGTCTTTGATTCTACAGAGGCAGAACAGTTAGGGGG*A*G |
| *Nutm2*–*Ywhae* t(13:11) | A*C*TGGCAGTCCAGAGTTCCACCTTCCCACTCAGGGAAGGCTTCTCCTTTA<br>TGAGAACTTGCTTGGTCAGCTTCTGAAAGCTCACTAGGTCATCAAGTA*T*T |
| *Adamts20*–*K18N* (Left) | C*G*TACTCGGGCCGGTCACAGAGCCGGGCTGTGCTCTTGATTCCACCTCCG<br>GGAGGGGATAATCAGTCAGGTGCCTCGGCTCAGGTTTCTCGGGTCAGC*T*T |
| *Adamts20*–*K18N* (Right) | G*A*TGGCGAGTGGGGACCATGGGGACCCTACAGCTCGTGTTCAAGGACATG<br>CGTATACACAGGGCTTCGCAGTTCCCAGGGCTCTTACGCATTTGATCC*T*C |

### Table S3: Primers used in the present study.

| Target loci | Genomic location (mm10/GRCm38) | Sequences (5′–3′) |
|---|---|---|
| **Genotyping primers** | | |
| Hmga2-1st-F | chr10: 120415727 to 120415748 | CCCCAGGGAAGGTAAATAATGT |
| Hmga2-1st-R | chr10: 120416637 to 120416658 | TTGCTCTGGACAACATTCATTC |
| Hmga2-2nd-F | chr10: 120416078 to 120416099 | GTCTGCCATGATGTTTGTTAGC |
| Hmga2-2nd-R | chr10: 120416271 to 120416292 | AAGTGTTGAAGAGCAGAAAGGC |
| Wif1-1st-F | chr10: 121049111 to 121049132 | TGACCCTCCACCATTAAAATTC |
| Wif1-1st-R | chr10: 121049971 to 121049992 | AGATCATTGCTGGGAAGAAGAG |
| Wif1-2nd-F | chr10: 121049423 to 121049444 | GTTCAGTAGCTGGAGGAGGATG |
| Wif1-2nd-R | chr10: 121049610 to 121049631 | GGCTTTTTATGAAAGGCAACTG |
| Rassf3-1st-F | chr10: 121431397 to 121431418 | GAAGACAGAGACAAACCATGCC |
| Rassf3-1st-R | chr10: 121432300 to 121432321 | CTCTTTGTGGCACTTCTGTGTC |
| Rassf3-2nd-F | chr10: 121431721 to 121431742 | GAGATTGCTCAGGACAGAGTGA |
| Rassf3-2nd-R | chr10: 121431917 to 121431944 | TGTACTCATACAAAATTAATCACAAGGG |
| Eef1a1N-1st-F | chr9: 78476997 to 78477018 | ATCCATGTGGAGCCTTTATCTC |
| Eef1a1N-1st-R | chr9: 78477804 to 78477825 | GTTAGCCTGACTTTGGCAGAAC |
| Eef1a1N-2nd-F | chr9: 78477215 to 78477236 | GCACTTAGCTTTCACGTGTTTG |
| Eef1a1N-2nd-R | chr9: 78477444 to 78477465 | TCAAGACCTACCTGGGAAATTG |
| Atf4N-1st-F | chr15: 80258094 to 80258115 | CCATTGGGTTTGCTAATCTAGG |
| Atf4N-1st-R | chr15: 80258876 to 80258897 | TGCTACTAGAGGTCTTGGGGAC |
| Atf4N-2nd-F | chr15: 80258497 to 80258518 | TGTGGGTAGGATGATACAGCAG |
| Atf4N-2nd-R | chr15: 80258705 to 80258726 | CCAAACCTGAGCTGGTCTATTT |
| Ypel4N-1st-F | chr2: 84738389 to 84738410 | ATCTAATCTTTGCCATTGGGTG |
| Ypel4N-1st-R | chr2: 84739219 to 84739240 | AACAACGCCTGGCAATTAATAC |
| Ypel4N-2nd-F | chr2: 84738693 to 84738714 | TTGTACTGTCCTCCGTATGCAC |
| Ypel4N-2nd-R | chr2: 84738873 to 84738894 | CCAGGGGACTGAAGACACTTAG |
| In(6)1J-left-1st-F | chr6: 63000428-63000449 | TCCCCAAAACATCACAATACAA |
| In(6)1J-left-1st-R | chr6: 63001303 to 63001324 | GATGGGGTTGACATTCCATAGT |
| In(6)1J-left-2nd-F | chr6: 63000607 to 63000628 | CCCTCCAGATTTTATCCCTCTC |
| In(6)1J-left-2nd-R | chr6: 63001001 to 63001022 | GCAGCTAAAAGAGTCAGCTTCA |
| In(6)1J-right-1st-F | chr6: 120826810 to 120826831 | AGTTAGTCCATGTTGTGGGCTT |
| In(6)1J-right-1st-F | chr6: 120827732 to 120827753 | ATACTCATACACAGGCACGCAC |
| In(6)1J-right-1st-F | chr6: 120826900 to 120826921 | TCACTGCAGAAATTCTTGGAAA |
| In(6)1J-right-1st-F | chr6: 120827428 to 120827449 | TCGTAATAAGGGCATTTCACCT |
| Ncoa2-1st-F | chr1: 13160923 to 13160944 | CTGGGAGTTCCAAACTATCAGC |
| Ncoa2-1st-R | chr1: 13161602 to 13161623 | GAAGTGAGTCCTGAGACAAGGG |
| Ncoa2-2nd-F | chr1: 13161102 to 13161123 | CACACACACAGCACTTCAACAC |
| Ncoa2-2nd-R | chr1: 13161429 to 13161450 | CACTTAGAATCCTGAGCCTTGG |
| Greb1-1st-F | chr12: 16733972 to 16733993 | AGCCGTCACAAACTTCTCTTTC |
| Greb1-1st-R | chr12: 16734652 to 16734673 | GGTTGCCCATCACTGTTAGAAT |
| Greb1-2nd-F | chr12: 16734188 to 16734209 | CAGTAACATCAGCTGGCACTCT |
| Greb1-2nd-R | chr12: 16734562 to 16734583 | ATCTGGGGAGACAGATGCTTTA |
| Ywhae-1st-F | chr11: 75760505 to 75760526 | GTCTTGCCTGCGTATATGATTG |
| Ywhae-1st-R | chr11: 75761127 to 75761148 | AATTCGAGACAGGGTTTCTCTG |
| Ywhae-2nd-F | chr11: 75760679 to 75760700 | TAGCCCAAGTGTGGGTTTTATT |
| Ywhae-2nd-R | chr11: 75761007 to 75761028 | TTCTCCTTAAGGCTGACAGAGG |
| Nutm2-1st-F | chr13: 50467549 to 50467570 | GTGTGGAGAGAGCTCTGGACTT |
| Nutm2-1st-R | chr13: 50468200 to 50468221 | AACTGAAACCAATAGCACTCCG |
| Nutm2-2nd-F | chr13: 50467737 to 50467758 | TGGTCCAGAGAAGGATCTTAGG |
| Nutm2-2nd-R | chr13: 50468110 to 50468131 | AGGAAACTGCAGAGATAGTGCC |
| Adamts20-1st-F | chr15: 94347286 to 94347307 | AGGAGGGACATCAGGTTACAGA |
| Adamts20-1st-R | chr15: 94348147 to 94348168 | TGTGATGACCACTGCATTATGA |
| Adamts20-2nd-F | chr15: 94347562 to 94347583 | TCTCTGAAACTCGCAGACTGAC |
| Adamts20-2nd-R | chr15: 94347823 to 94347844 | TTCCTGTGTGTGTGCTTCTTCT |
| K18N-1st-F | chr15: 102034486 to 102034507 | CCTTTGTTCCGTTGATCATGTA |
| K18N-1st-R | chr15: 102035201 to 102035222 | GAAACTGGAGGGTCACAGGTAG |
| K18N-2nd-F | chr15: 102034680 to 102034700 | GGCATCAAATGTGTCTTCTCA |
| K18N-2nd-R | chr15: 102034945 to 102034966 | TGATGTCTGTGGCCTTTACTGT |
